# On the use of inhibitors of 4-hydroxyphenylpyruvate dioxygenase as a vector-selective insecticide in the control of mosquitoes

**DOI:** 10.1101/669747

**Authors:** Marlon A. V. Ramirez, Marcos Sterkel, Ademir de Jesus Martins, José Bento Pereira Lima, Pedro L. Oliveira

**Affiliations:** Laboratório de Bioquímica de Artrópodes Hematófagos, Instituto de Bioquímica Médica Leopoldo de Meis, Universidade Federal do Rio de Janeiro, Rio de Janeiro, RJ, Brazil; Centro Regional de Estudios Genómicos, Universidad Nacional de La Plata (CREG-UNLP), Argentina; Laboratorio de Fisiologia e Controle de Artrópodes Vetores, Instituto Oswaldo Cruz, FIOCRUZ, Rio de Janeiro, RJ, Brasil; Laboratório de Entomologia, Instituto de Biologia do Exército, Rio de Janeiro, RJ, Brasil; Instituto Nacional de Ciencia e Tecnologia em Entomologia Molecular (INCT-EM), Brazil

## Abstract

Blood-sucking insects incorporate many times their body weight of blood in a single meal. As proteins are the major component of vertebrate blood, its digestion in the gut of hematophagous insects generates extremely high concentrations of free amino acids. Previous reports showed that the tyrosine degradation pathway plays an essential role in adapting these animals to blood feeding. Inhibiting 4-hydroxyphenylpyruvate dioxygenase (HPPD), the rate-limiting step of tyrosine degradation, results in the death of insects after a blood meal. Therefore, it was suggested that compounds that block the catabolism of tyrosine could act selectively on blood-feeding insects. Here we have evaluated the toxicity against mosquitoes of three HPPD inhibitors currently used as herbicides and in human health. Among the compounds tested, nitisinone (NTBC) proved to be more potent than mesotrione (MES) and isoxaflutole (IFT) in *Aedes aegypti*. NTBC was lethal to *Ae. aegypti* in artificial feeding assays (LD50: 4.36 µM), as well as in topical application (LD50: 0.0033 nmol/mosquito). NTBC was also lethal to *Ae. aegypti* populations that were resistant to neurotoxic insecticides, and it was lethal to other mosquito species (*Anopheles* and *Culex*). Therefore, HPPD inhibitors, particularly NTBC, represent promising new drugs for mosquito control. Since they only affect blood-feeding organisms, they would represent a safer and more environmentally friendly alternative to conventional neurotoxic insecticides.

**Author Summary:** The control of mosquitoes has been pursued in the last decades by the use of neurotoxic insecticides to prevent the spreading of dengue, zika and malaria, among other diseases. However, the selection and propagation of different mechanisms of resistance hinder the success of these compounds. New methodologies are needed for their control. Hematophagous arthropods, including mosquitoes, ingest quantities of blood that represent many times their body weight in a single meal, releasing huge amounts of amino acids during digestion. Recent studies showed that inhibition of the tyrosine catabolism pathway could be a new selective target for vector control. Thus we tested three different inhibitors of the second enzyme in the tyrosine degradation pathway as tools for mosquito control. Results showed that Nitisinone (NTBC), an inhibitor used in medicine, was the most potent of them. NTBC was lethal to Aedes aegypti when it was administered together with the blood meal and when it was topically applied. It also caused the death of Anopheles aquasalis and Culex quinquefasciatus mosquitoes, as well as field-collected Aedes populations resistant to neurotoxic insecticides, indicating that there is no cross-resistance. We discuss the possible use of NTBC as a new insecticide.

## Introduction

Mosquitoes are important vectors for pathogens that cause diseases such as malaria, lymphatic filariasis, yellow fever, dengue, chikungunya, Zika and West Nile fever. Mosquito-borne diseases are among the leading global public health menaces [1], and vector control by means of insecticides is crucial for the management of these diseases [2,3]. Conventional insecticides are applied in the stage of larva and adult. To reduce these immature forms in the case of *Ae. aegypti,* temephos (organophosphate) has been used for many years, as well as mechanical control to eliminate standing water [4]. For adult control, malathion (organophosphate) and pyrethroids are mainly recommended by World Health Organization (WHO). In the case of *Anopheles* mosquitoes, insecticide-treated nets (ITNs) and indoor residual spraying (IRS) are used as preventive strategies [5]. The main classes of insecticides are organochlorines, organophosphates, carbamates, and pyrethroids, all of them neurotoxic [6]. However, their extensive use has led to the development of resistance to these insecticides [7,8], representing a problem for mosquito control. This is particularly true for arbovirus transmitted by *Ae. aegypti*, exemplified by the recent global Zika outbreak [9], making the search for alternative methods for mosquito control a high priority of the global public health agenda [10].

Female mosquitoes need to feed on blood for the maturation of their eggs. Since 85% of vertebrate blood dry weight is protein, its digestion generates high concentrations of amino acids in the gut [11]. Although amino acids are considered essential nutrients, many human genetic diseases are caused by defective degradation of amino acids, leading to hyperaminoacidemias and the formation of toxic metabolites [12]. In the hematophagous “kissing bug” *Rhodnius prolixus,* a high level of expression of enzymes related to tyrosine degradation is found in the midgut [13]. Silencing of 4-hydroxyphenylpyruvate dioxygenase (HPPD), the enzyme that catalyzes the second (and rate-limiting) step of the tyrosine degradation pathway, led to insect death after a blood meal. Chemical inhibitors of HPPD caused the death of hematophagous arthropods (kissing bug, mosquitoes and ticks) but were not toxic to non-hematophagous insects [14]. Therefore, the degradation of free tyrosine formed in excess during digestion of a blood meal is an essential trait in the adaptation to a hematophagous way of life [11]. The key role of this pathway in the evolution of blood-feeding organisms led us to identify tyrosine degradation as a potential target in the development of novel alternative insecticides that would therefore be selective for these animals.

In plants, functional HPPD is required for the synthesis of plastoquinone and tocopherol, which are essential for the plant to survive. Therefore, HPPD has been identified as one of the most promising targets for the development of new herbicides, and thousands of HPPD inhibitors have been synthesized [15]. HPPD inhibitors are classified into three main chemical families: triketones, diketonitriles and pyrazolones [16,17]. Triketones can be natural compounds such as leptospermone, or synthetic, such as mesotrione (MES), nitisinone (NTBC) and many others [18]. MES is used as a herbicide (Callisto®, Syngenta) [19]. NTBC, on the other hand, is approved for therapeutic use in humans to treat hereditary tyrosinemia type 1 (HT-1) since 1994 (Orfadin®) [20] and its potential for use in patients with alkaptonuria is under investigation [21,22]. In the diketonitriles group, isoxaflutole (IFT) is also used as a herbicide (Balance® and Merlin®, Bayer) [23]. To test the possibility of using HPPD inhibitors as selective insecticides for disease vectors, here we investigated the toxicity of three inhibitors of HPPD (NTBC, MES and IFT) towards mosquitoes, especially against *Ae. aegypti*, evaluating different doses and possible modes of application. We also tested NTBC toxicity to *Ae. aegypti* populations resistant to pyrethroids and organophosphates, as well as other species of mosquitoes, *Culex quinquefasciatus* and *Anopheles aquasalis*. Our results support the use of HPPD inhibitors, particularly NTBC, as new insecticides for mosquito control.

## Methods

### Ethics statement

All experiments were conducted according to the guidelines of the institutional care and use committee (Committee for Evaluation of Animal Use for Research from the Federal University of Rio de Janeiro, CAUAP-UFRJ), which is based on the NIH Guide for the Care and Use of Laboratory Animals (ISBN 0-309-05377-3). The protocols used here were approved by CAUAP-UFRJ under registry #IBQM155/13.

### Mosquito rearing

*Ae. aegypti* Red Eye strain were maintained in the insectary of the Federal University of Rio de Janeiro (UFRJ), Brazil. The insecticide-resistant *Ae. aegypti* populations, originally collected with ovitraps from different Brazilian cities: Santarém (Pará state), Nova Iguaçu (Rio de Janeiro state), and Oiapoque (Amapá state), were reared in the insectary at FIOCRUZ-RJ. A Rock-kdr strain (or R2R2 strain) that presents point mutations in the voltage-dependent sodium channel associated with pyrethroid resistance was also maintained in the FIOCRUZ-RJ insectary [24,25]. The *Ae. aegypti* Rockefeller strain (also maintained at FIOCRUZ-RJ), which is commonly used as a standard for insecticide susceptibility assays, was used here as a reference strain, in order to allow better comparison with literature data [26]. RR50 (Resistance rate 50: LD50 of strain or population studied/LD50 of Rockefeller strain) for deltamethrin was: Santarém population = 30.4, Nova Iguaçu population = 25.4, Oiapoque population = 143.9 [27–30]. In the case of Rock-kdr, a knockdown time assay with deltamethrin revealed that the time necessary to knockdown 95% of the Rock-kdr lineage was 6.7x longer than the time found for the Rockefeller susceptible strain [25].

The larvae were fed with powdered cat food (Friskies^®^, Nestlé Purina PetCare). Adult mosquitoes were kept in cages and fed with a 10% sucrose solution. All mosquitoes were reared at 26°C, in 70-80% relative humidity and a photoperiod of 12h light:12h dark.

A colony of *An. aquasalis* was established in 1995 using specimens collected in the municipality of Guapimirim, Rio de Janeiro, and reared in the insectary of FIOCRUZ-RJ. The larvae were reared on a diet of fish food (Tetra Marine Large Flakes, Tetra GmH) in containers containing dechlorinated water at a concentration of 0.2 % NaCl (w/v). Adult mosquitoes were provided with 10 % sucrose *ad libitum* under a regimen of photoperiod, temperature and humidity similar to that of *Ae. aegypti* [31].

*Cx. quinquefasciatus* were reared at Instituto de Biologia do Exército (IBEX), Rio de Janeiro. The larvae were fed cat food (Friskies®, Nestlé Purina PetCare). To encourage copulation, adults were housed in a dark room, because this mosquito feeds on blood at night. Temperature and humidity were similar to the other mosquitoes.

### Topical application assays

NTBC, MES or IFT (Sigma Chemical Co.) were diluted in acetone just before each experiment. After feeding on blood, mosquitoes were cold-anesthetized and placed in a glass petri dish on ice. Then, a volume of 0.5 μl of the HPPD inhibitor solution was topically applied with a micropipette on the abdomen of the insect. Survival was evaluated every 24 h for a week. Controls received only acetone.

### Artificial feeding assays

Rabbit blood was collected with a sterile syringe containing heparin at a ratio of 1 μl heparin stock/ml blood (heparin stock was 5000 IU/ml). Stock solutions of HPPD inhibitors in PBS (NaCl 0.15 M, Na phosphate 10 mM, pH 7.0) were diluted in PBS and mixed 1:9 (v/v) with heparinized blood to obtain final concentrations used to feed mosquitoes, as indicated in figure legends. Controls received PBS in blood (1:9, v/v). Mosquitoes of 3-5 days post-emergence were fed in an artificial feeding apparatus where food was offered through a membrane of Parafilm M®. The temperature of the blood meal was maintained with a circulating water bath, adjusted at 37-38°C [32]. The maximum feeding time was 30 minutes. Only fully engorged mosquitoes were used. Survival was evaluated every 24 h for a week.

### Survival experiments, statistical analysis

Mosquitoes were offered a 10% sucrose solution *ad libitum* and were considered dead if they could no longer stand. Statistical analysis and design of graphs were performed using Prism 6.0 software (GraphPad Software, San Diego, CA). At least two independent experiments were performed for each experimental condition (each with its respective control group). The Kaplan-Meier survival curve analysis in the Prism software (the log rank test) was used to evaluate significant differences between experimental and control groups. LD50 was determined using a non-linear regression to fit log[inhibitor] vs. normalized response (Variable slope). Two-way analysis of variance and Tukey’s multiple comparisons test were carried out to compare the field populations and laboratory strains with Rockefeller strain.

## Results

We examined three chemical inhibitors of HPPD for their effects on the survival of *Ae. aegypti* (Red Eye strain): two of them are marketed as herbicides (MES and IFT), and one is used in medicine for the treatment of tyrosinemia type I (NTBC). MES and NTBC are triketones while IFT belongs to the diketonitrile family. Inhibitors from the family of pyrazoles were not evaluated. The three inhibitors were administered by artificial feeding and topical application. NTBC was the most potent of the three inhibitors in the artificial feeding trials, presenting an LD50 (the dose that kills 50% of mosquitoes) of 4.36 µM, while MES presented an LD50 of 324 µM (Fig. 1, Table 1). IFT showed no lethal effects in any of the concentrations tested. Thus, NTBC was about 74 times more potent than MES when it was co-administered along with the blood meal.

**Figure 1.**
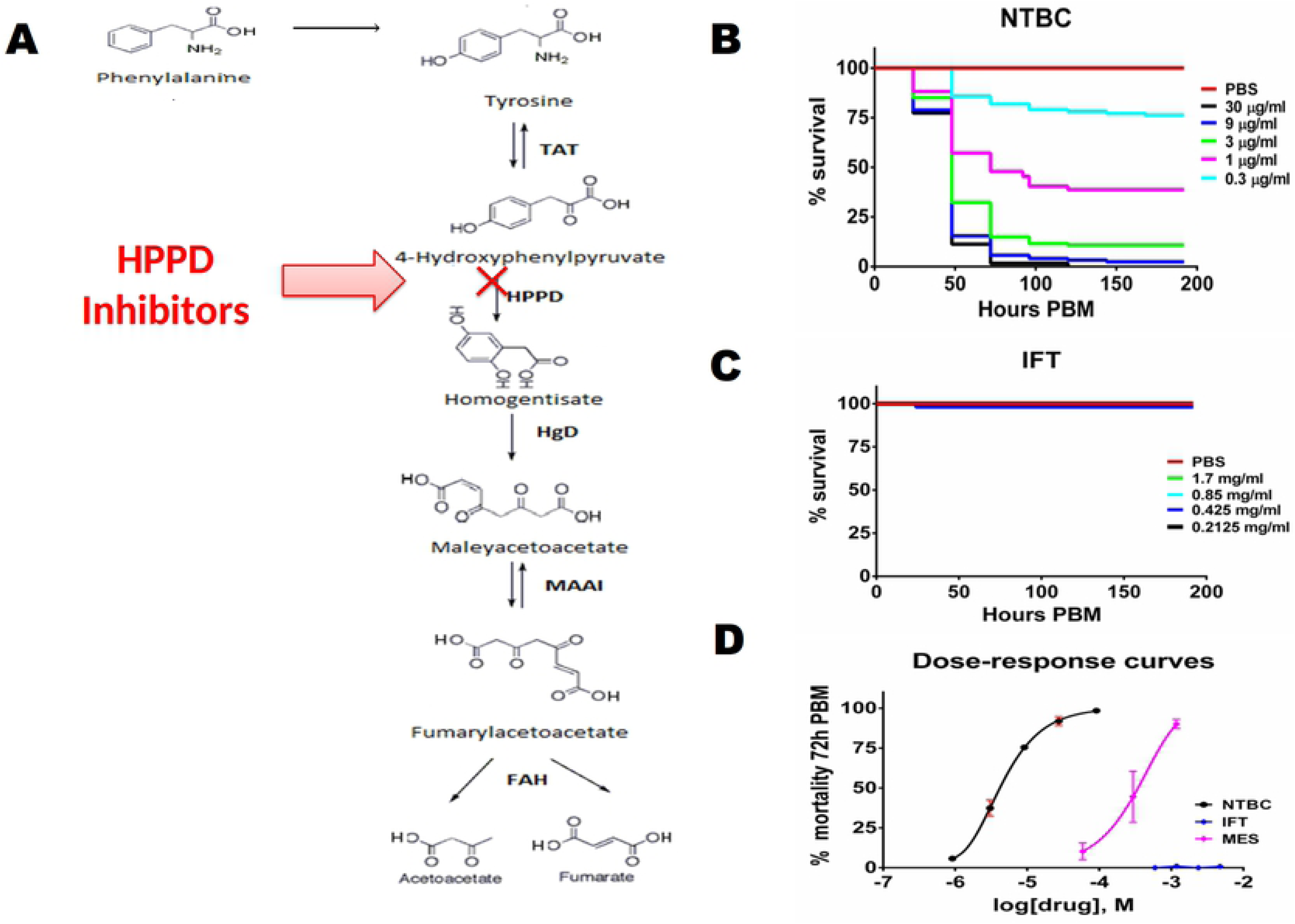
Ingestion of HPPD inhibitors with the blood meal decreases the survival of *Ae. aegypti* (Red Eye strain). **(A)** Tyrosine catabolism pathway. TAT: tyrosine aminotransferase; HPPD: 4-hydroxyphenylpyruvate dioxygenase; HgD: homogentisate 1, 2 dioxygenase; MAAI: maleylacetoacetate isomerase; FAH: fumarylacetoacetase. **(B)** Survival rates of *Ae. aegypti* (Red Eye strain) fed with rabbit blood supplemented with NTBC. PBM: Post-blood meal. Control group was fed with blood plus PBS (9:1; v/v). **(C)** Survival rates of *Ae. aegypti* (Red Eye strain) fed with rabbit blood supplemented with IFT **(D)** Dose-response curves at 72 h PBM. MES data were taken from Sterkel *et al.* (2016). Four (panel B) and two (panel C) independent experiments were performed respectively, each with n =10–36 insects per experimental group. Panels B and C are plotted as Kaplan-Meier survival curves.

**Table 1.**
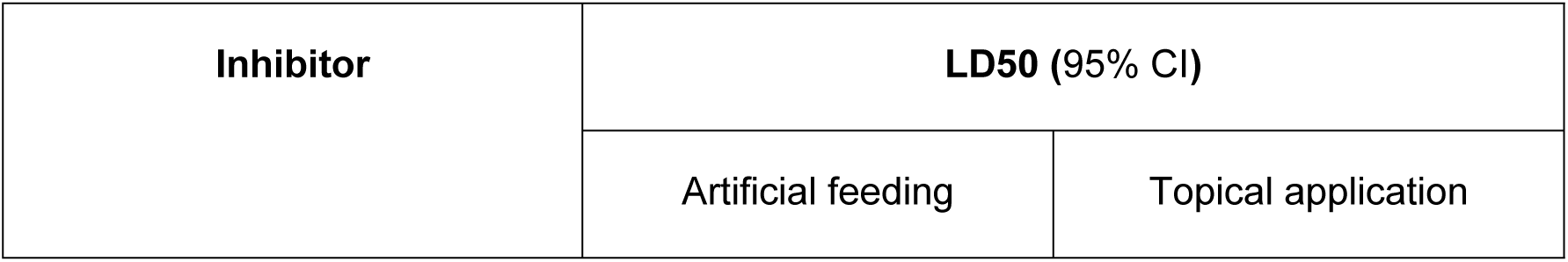
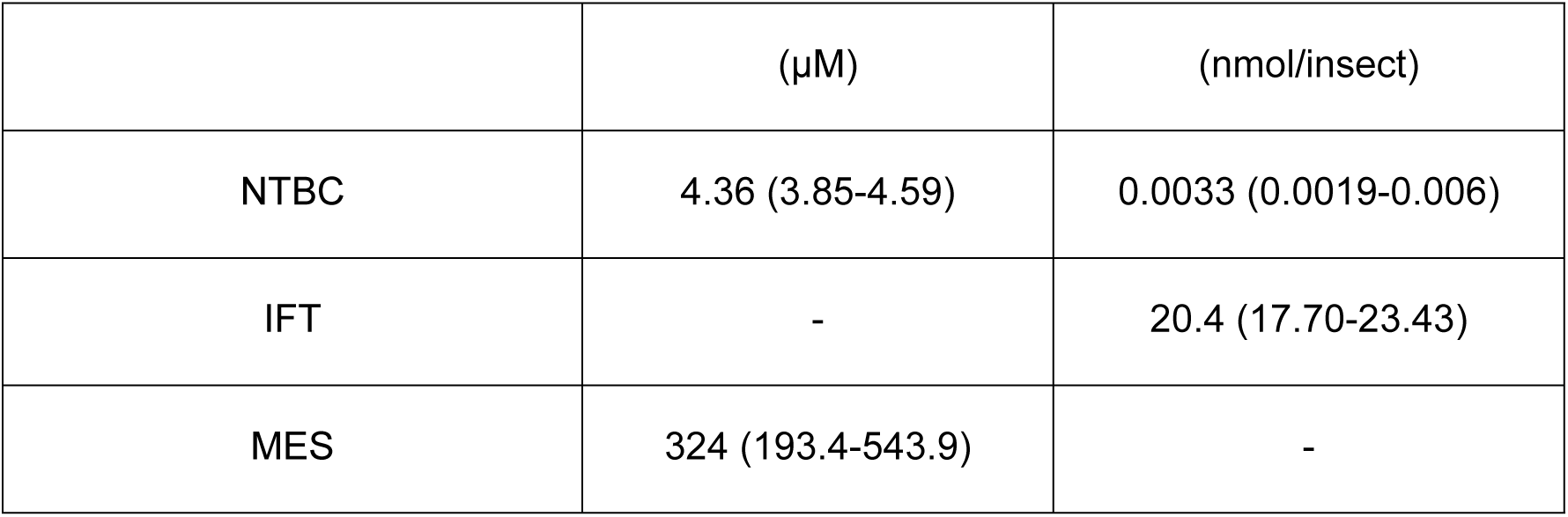
Toxicity of HPPD inhibitors to *Ae. aegypti* (Red Eye strain) using artificial feeding and topical application - LD50 were calculated from data in Figures 1 and 2. Data shown are mean ± 95% Confidence Interval (CI).

| Inhibitor | LD<sub>50</sub> (95% CI) |  |
| --- | --- | --- |
|  | Artificial feeding | Topical application |

| | (μM) | (nmol/insect) |
| --- | --- | --- |
| NTBC | 4.36 (3.85-4.59) | 0.0033 (0.0019-0.006) |
| IFT | - | 20.4 (17.70-23.43) |
| MES | 324 (193.4-543.9) | - |

NTBC proved to be more potent than MES and IFT also in the topical application assay (Figure 2), presenting an LD50 = 0.0033 nmol/insect (1.1 ng/insect). However, in contrast to the artificial feeding assay, IFT also was lethal, presenting an LD50 = 20.4 nmol/insect (7320 ng/insect) (Table 1). These results demonstrated that NTBC was around 6182 times more potent than IFT. On the other hand, MES did not cause mortality in mosquitoes when applied topically, suggesting that this compound was not able to traverse the cuticle.

**Figure 2.**
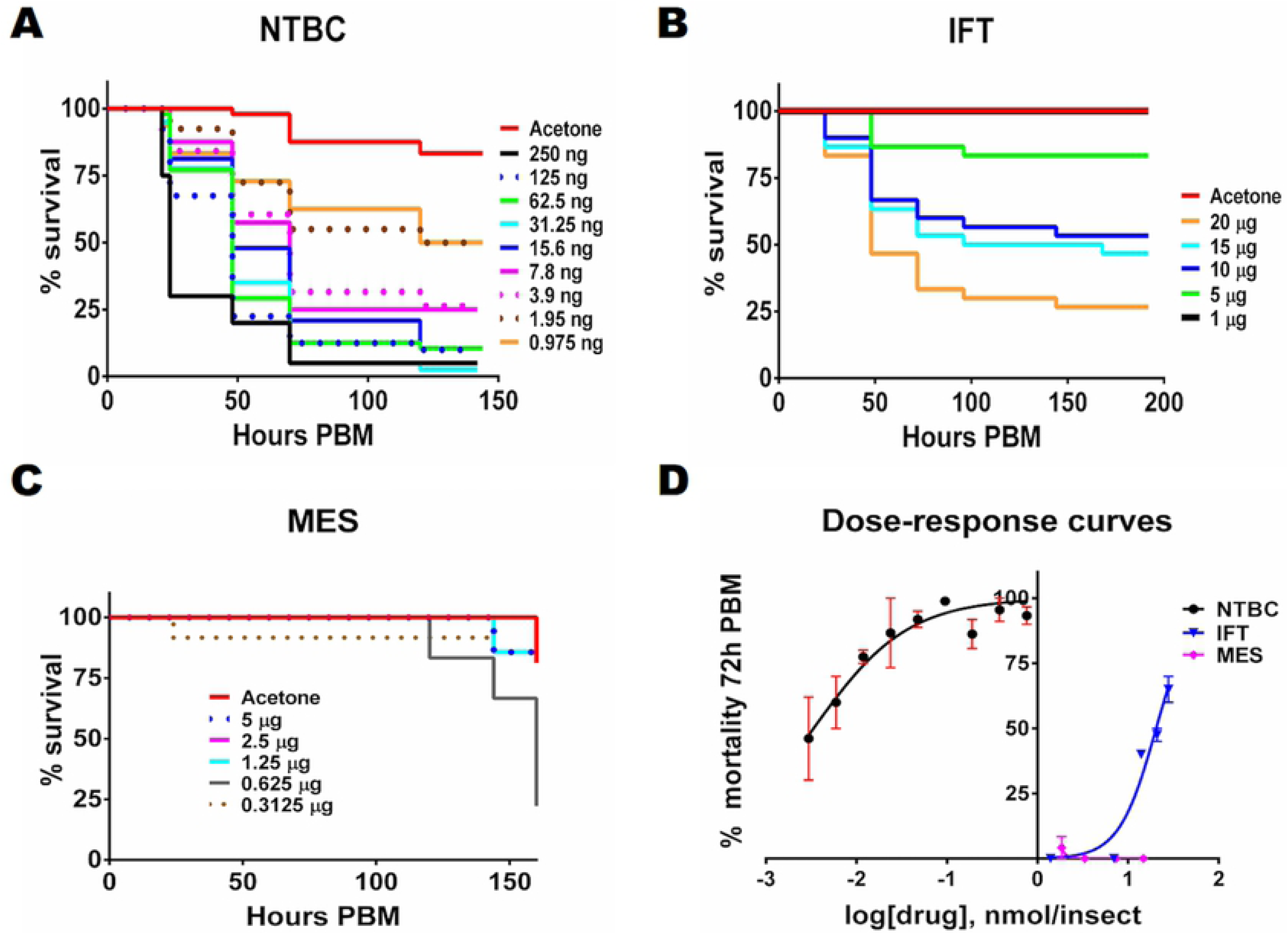
Topical application of HPPD inhibitors effect on survival of *Ae. aegypti* (Red Eye strain) after a blood meal. Topical application of **(A)** NTBC or **(B)** IFT on the abdomen causes mosquitoes death. **(C)** MES had very little effect. **(D)** Dose-response curves recorded at 72 h PBM. Panels A, B and C are plotted as Kaplan-Meier survival curves. Three (A) and two (B, C) independent experiments were performed, respectively, each with n =12–35 insects per experimental group. The drugs (dissolved in 0.5 µl acetone) were applied on the abdomen immediately after the blood meal (time 0 PBM). Controls received only acetone (0.5 µl).

The effect of NTBC on other mosquito species was evaluated in trials of artificial feeding. *Cx. quinquefasciatus* and *An. aquasalis* died when fed with blood containing concentrations of NTBC similar to those that killed *Ae. aegypti* (Red Eye and Rockefeller strains) (Fig. 3 and Table 2). These results show that NTBC can be used not only for the control of *Aedes* populations, but also for the control of other mosquitoes that transmit pathogens.

**Figure 3.**
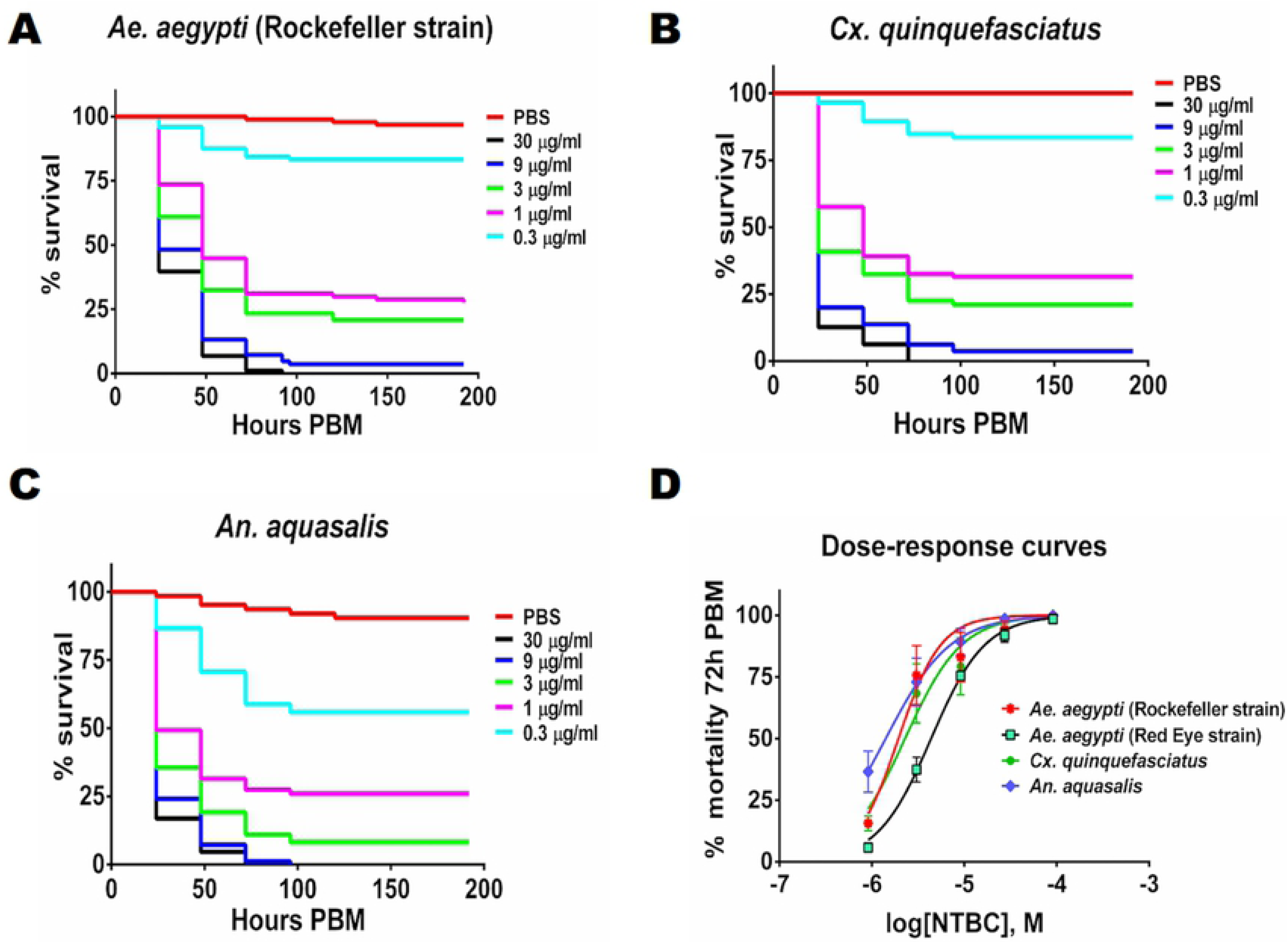
Ingestion of NTBC is lethal to other mosquito species. **(A)** *Ae. aegypti* (Rockefeller strain) were fed on rabbit blood plus NTBC. **(B)** *Cx. quinquefasciatus* were fed rabbit blood plus NTBC. **(C)** *An. aquasalis* were fed rabbit blood plus NTBC. **(D)** Dose-response curves at 72 h PBM. Panels A, B and C are plotted as Kaplan-Meier survival curves. 3 independent experiments were performed, each with n =10–37 insects per experimental group.

**Table 2.** Toxicity of NTBC to different mosquito species using artificial feeding - LD50 were calculated from data in Figure 3. Data shown are mean ± 95% Confidence Interval (CI).

| Species of mosquito | NTBC LD50 (95% CI) |  |
| --- | --- | --- |
|  | µM | µg/ml |
| *Ae. aegypti* (Rockefeller strain) | 1.94 (1.42-2.65) | 0.64 µg/ml |
| *Cx. quinquefasciatus* | 2.28 (1.56-3.34) | 0.75 µg/ml |
| *An. aquasalis* | 1.41 (1.03-1.93) | 0.46 µg/ml |

Next, we studied the effect of NTBC on field-collected populations of *Ae. aegypti* that showed high levels of resistance to organophosphates and pyrethroids. NTBC was also tested on the laboratory Rock-kdr strain, which carries a mutation in the voltage-dependent sodium channel, the target site of pyrethroids and organochlorines (DDT). When these populations were fed with NTBC-supplemented blood, they presented LD50 values similar to the control (Rockefeller strain) (Fig. 4 and Table 3). Thus neither field-collected *Ae. aegypti* populations nor the Rock-kdr strain showed cross-resistance to NTBC when orally administered along with a blood meal.

**Figure 4.**
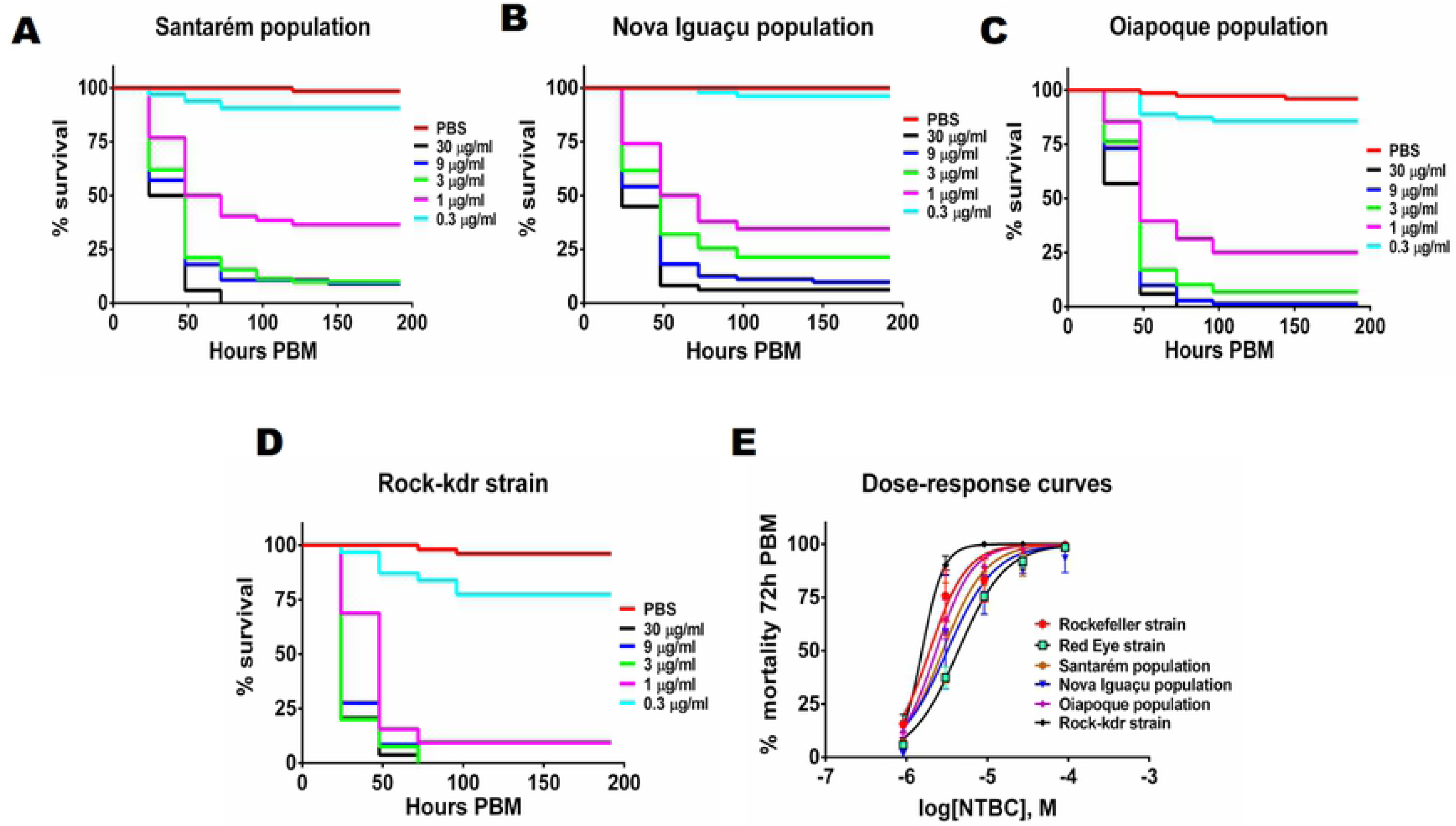
Insecticide-resistant *Ae. aegypti* populations do not show cross-resistance towards NTBC ingested with a blood meal. Different *Ae. aegypti* populations were fed with blood plus NTBC. **(A)** Santarém population, **(B)** Nova Iguaçu population, **(C)** Oiapoque population, **(D)** Rock-kdr strain. **(E)** Dose-response curves at 72 h PBM. Panels A-D are plotted as Kaplan-Meier survival curves. Three (A, B, C) and two (D) independent experiments were performed, respectively, each with n =10–35 insects per experimental group.

**Table 3.** Toxicity of NTBC to insecticide-resistant *Ae. aegypti* populations - LD50 were calculated from data in Figures 4 and 5. ***= p<0.001. Data shown are mean ± 95% Confidence Interval (CI).

| ***Ae. aegypti* strains or populations** | **NTBC LD50 (95% CI)** |  |
| --- | --- | --- |
|  | Artificial feeding<br>(µM) | Topical application<br>(nmol/insect) |
| Rockefeller | 1.94 (1.42-2.65) | 0.0046 (0.0036-0.0060) |
| Santarém | 2.82 (1.79-4.41) | 0.0235 (0.017-0.03)*** |
| Nova Iguaçu | 3.20 (1.76-5.79) | 0.0255 (0.018-0.037)*** |
| Oiapoque | 2.33 (1.94-2.76) | 0.0181 (0.012-0.026)*** |
| Rock-kdr | 1.56 (1.39-1.75) | 0.0075 (0.0052-0.011) |

Finally, topical application trials were also carried out on the insecticide-resistant mosquitoes. In this case, unlike artificial feeding, there were significant differences between field-collected neurotoxic-resistant populations and controls (Rockefeller strain) (p<0.001). These populations were less susceptible to NTBC than controls, presenting RR50 values of 5.1, 5.5 and 3.9 for Santarém, Nova Iguaçu and Oiapoque populations, respectively. In contrast, there were no significant differences between the LD50 values calculated for Red Eye, Rock-kdr and Rockefeller strains (Figure 5 and Table 3). These results might be explained by a reduced penetration of this drug through the cuticle, since no major differences were observed in NTBC LD50 during artificial feeding trials.

**Figure 5.**
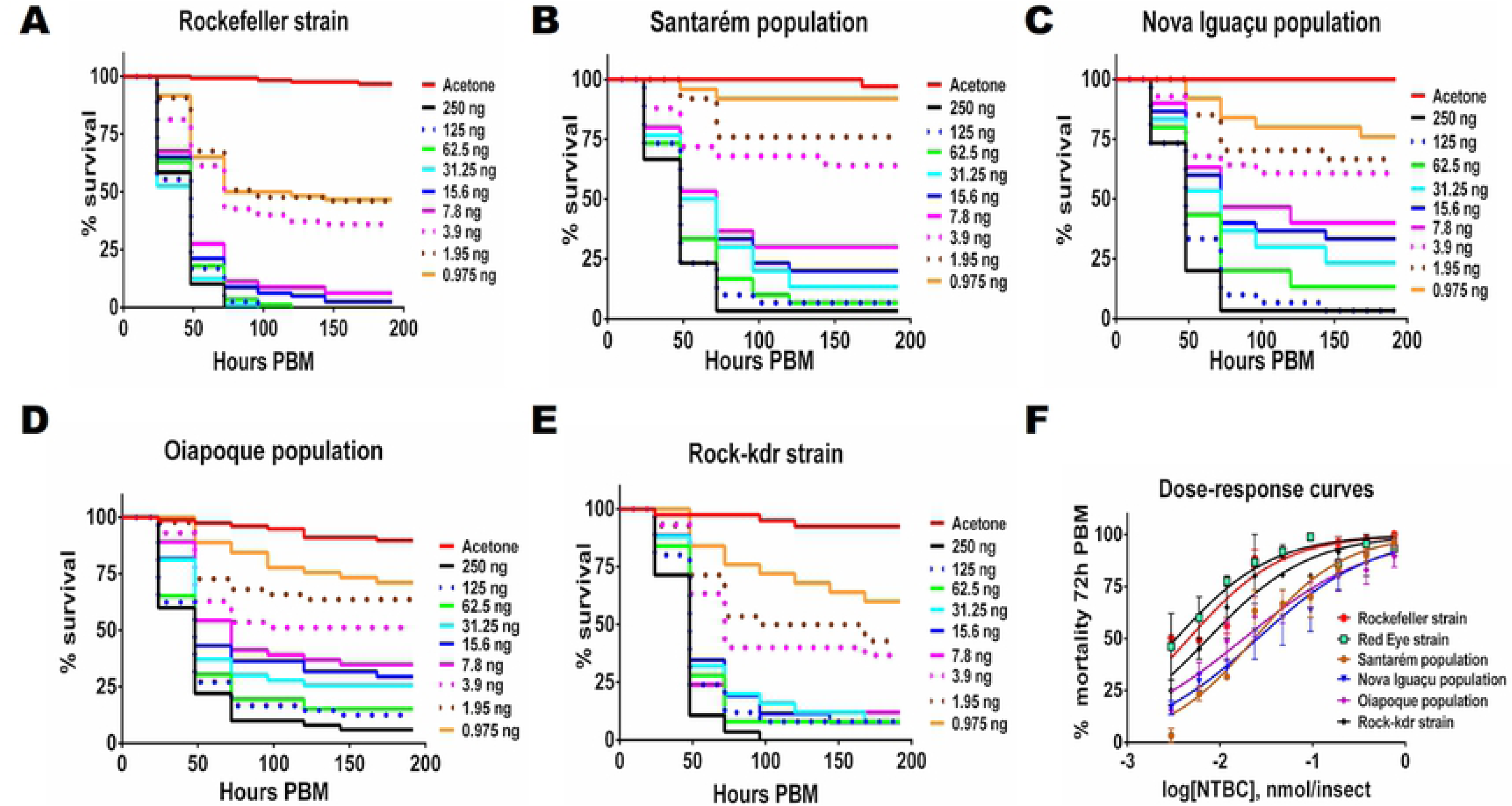
Insecticide-resistant *Ae. aegypti* populations show moderate resistance to NTBC applied topically after a blood meal. Topical application of NTBC caused death to *Ae. aegypti* **(A)** Rockefeller, **(B)** Santarém, (**C)** Nova Iguaçu, **(D)** Oiapoque; and **(E)** Rock-kdr strain. **(F)** Dose-response curves at 72 h PBM. Panels A, B, C, D and E are plotted as Kaplan-Meier survival curves. Four independent experiments for Rockefeller strain and Oiapoque population and two independent experiments for the other mosquito populations were performed, each with n =10–32 insects per experimental group. The drug (in 0.5 µl acetone) was applied on the abdomen immediately after a blood meal. Controls received 0.5 µl acetone. Differences in the LD50 between Santarém, Nova Iguaçu and Oiapoque populations (p<0.001) and Rockefeller strain were observed.

Toxicity of HPPD inhibitors toward hematophagous arthropods depends on the digestion of blood meal proteins, as previous data showed that sugar-fed female mosquitoes were not sensitive to mesotrione [14]. Using topical/contact applications of the drug under field conditions ensures that drug exposure does not occur at the same time as blood intake. Therefore, we decided to determine the maximum time interval between drug exposure and blood feeding that would still ensure lethality of the drug. For this, mosquitoes were fed on blood at different times (0 h, 24 h, 48 h, 72 h, 96 h) after the topical application of an LD95 of NTBC (15.6 ng/mosquito, the dose that kills 95% of the mosquitoes when it is applied immediately after a blood meal). Although a significant toxicity was retained for the first 24 h after drug application, the longer the time between blood feeding and topical application of NTBC, the lower the number of dead mosquitoes, suggesting progressive inactivation and/or excretion of the drug by the mosquito (Figure 6). In a second set of tests, mosquitoes were first fed on blood and then the topical application was made at different times PBM (0 h, 24 h, 48 h, 72 h) (Figure 7). Here, a similar time-dependent loss of efficacy is observed because by 48 h and later time points after the meal, there is little blood protein in the gut, and tyrosine has already been catabolized.

**Figure 6.**
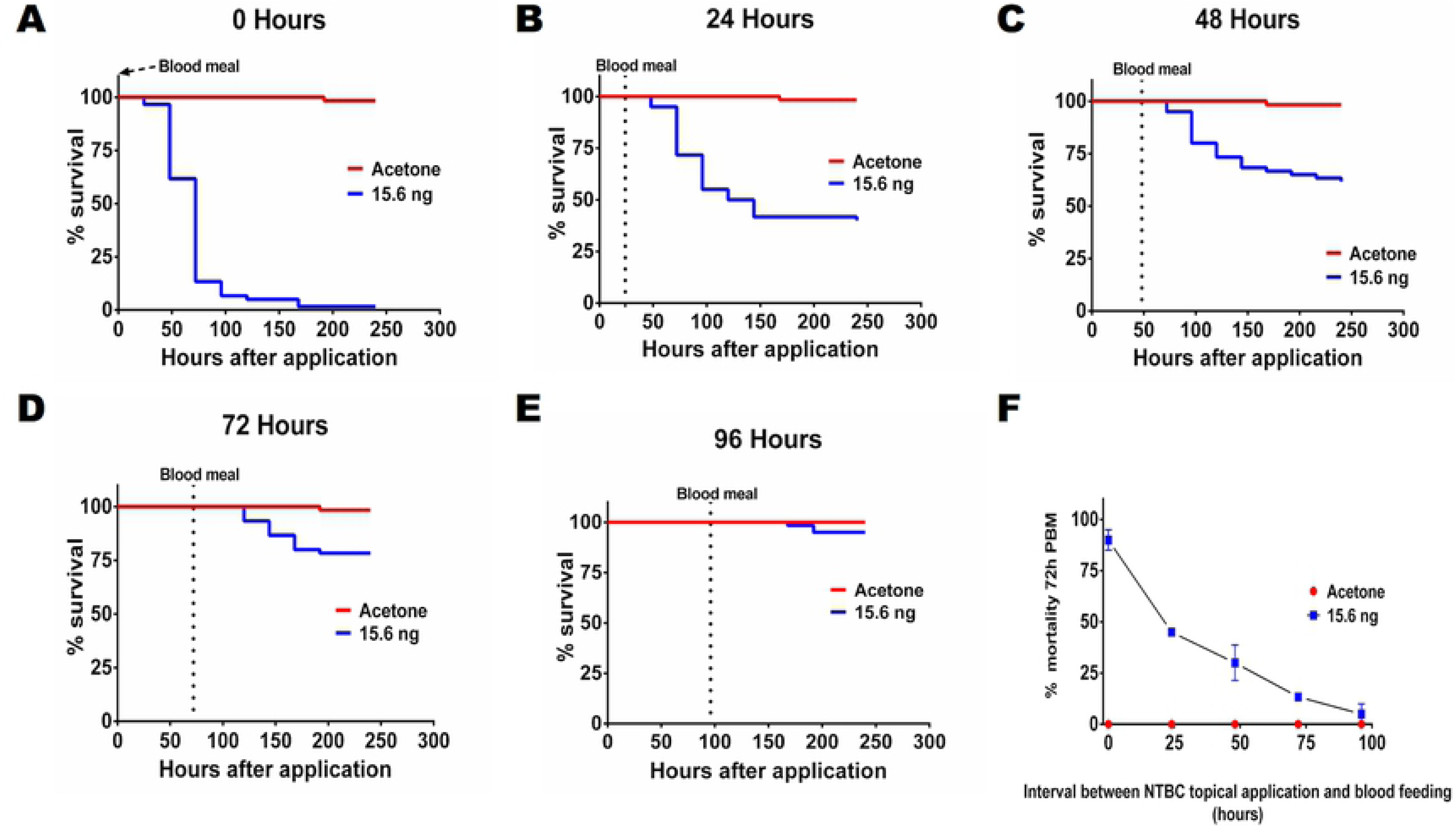
Blood meal administered at different times after topical application of NTBC. To evaluate persistence of toxicity from topically applied NTBC on *Ae. Aegypti* (Red Eye strain) the drug was applied at time 0 and the blood meal was administered at different times (dotted line): (**A**) 0 h, (**B**) 24 h, (**C**) 48 h, (**D**) 72 h, (**E**) 96 h. The results of A-E are summarized as % mortality observed at 72 h PBM in (**F)**. Panels A-E are plotted as Kaplan-Meier survival curves. Two independent experiments were performed, each with n =10–32 insects per experimental group. The LD95 (15.6 ng in 0.5 µl acetone) was applied on the abdomen of the mosquito at time 0. Acetone alone was applied to the control group at 0 h PBM.

**Figure 7.**
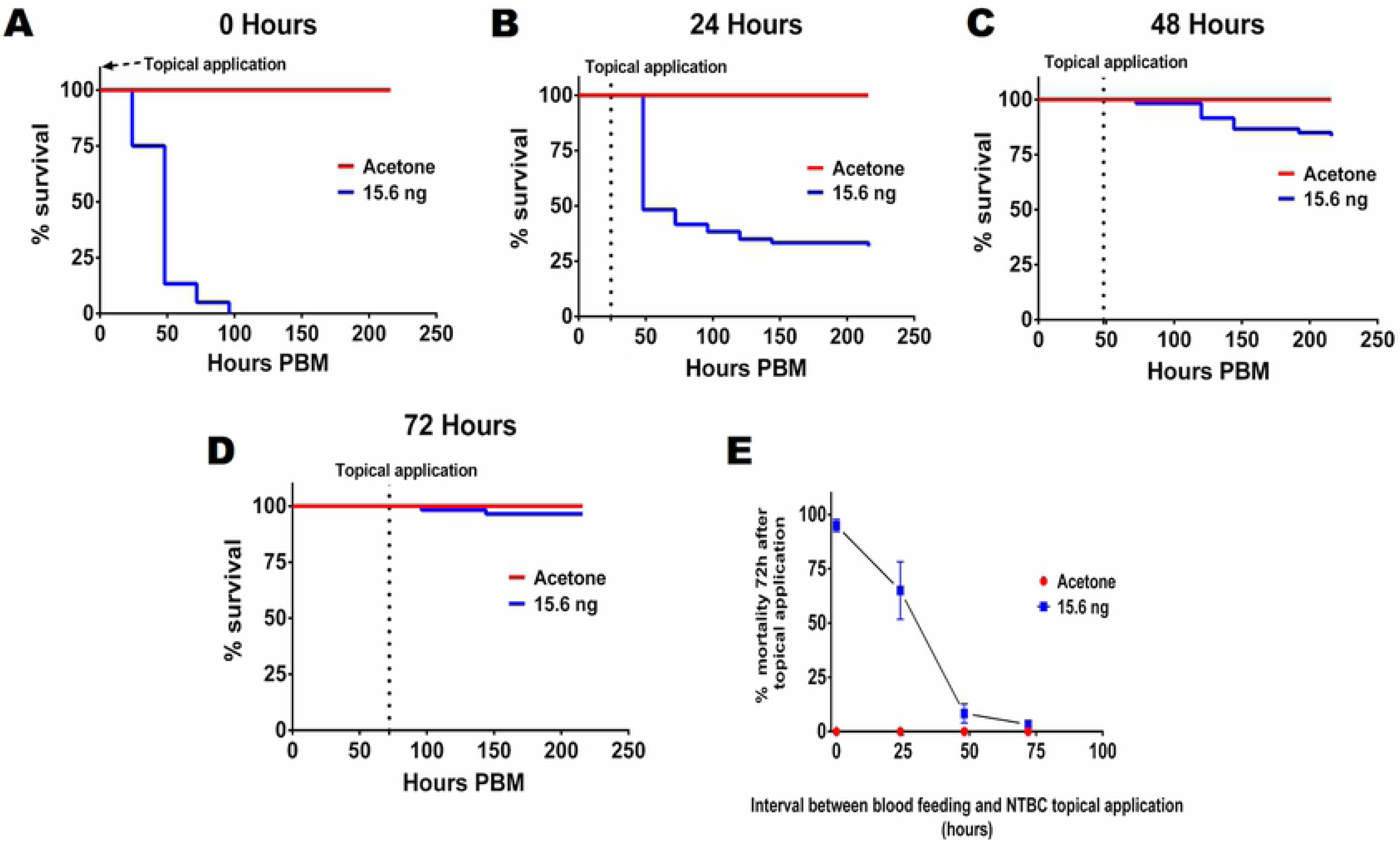
Topical application of NTBC at different times after PBM. Mosquitoes (Red Eye strain) were fed with blood meal at time 0 and topical application of NTBC was performed at different times (dotted line): **(A)** 0 h, **(B)** 24 h, **(C)** 48 h, **(D)** 72 h. The results of A-D are summarized as % mortality observed at 72 h after topical application **(E)**. Data from panels A to D were used to summarize the effect of time interval between NTBC application and blood meal on mortality observed at 72 h after topical application. Data in A-D are plotted as Kaplan-Meier survival curve. Two independent experiments were performed, each with n =10–32 insects per experimental group.

## Discussion and Conclusions

Hematophagy in arthropods is linked to a hyperproteic diet to an extent not found in other animals [11]. When the proteins in the blood are degraded, high levels of amino acids such as tyrosine accumulate in the digestive tract. The discovery that the capacity to degrade free tyrosine produced in excess during blood meal digestion is an essential trait in the physiology of blood-sucking arthropods that contributes to adapt these animals to hematophagy led us to propose the use of HPPD inhibitors as a new class of insecticides, selective for hematophagous animals [14]. In this study we evaluated the potential use of HPPD as a novel target for the control of mosquitoes, comparing different inhibitors and modes of application. Our results showed that neither MES nor IFT was very powerful in *Ae. aegypti* (Red Eye strain), while NTBC stood out for its potency and efficacy. This was consistent with the report of Sterkel *et al*. (2016), who found that feeding of *Ae. aegypti* with rabbit blood supplemented with MES decreased their survival, as did feeding them on mice treated with an orally applied therapeutic dose of NTBC [14]. However, differences in the results depending on the mode of administration provided relevant insights that can help further research on the use of HPPD inhibitors as vector-selective insecticides. NTBC and MES, but not IFT, were effective against *Ae. aegypti* when administered along with the blood meal (Figure 1), but topical application resulted in a different profile. Although much higher doses were required, NTBC and IFT applied topically, but not MES, were effective.

An open question concerns the molecular mechanism responsible for the differences between MES, IFT and NTBC in response to topical application and artificial feeding, knowing that these compounds act on the same enzyme, HPPD. Whereas MES and NTBC act directly on the HPPD enzyme[19,33], IFT undergoes a biotransformation to the diketonitrile derivative (DKN), and it is this compound that acts on the HPPD enzyme [23,34]. Specifically, the difference between MES and IFT in these two assays might be explained either by differential absorption or by differential metabolic modification of these compounds when administered by each route (cuticle and midgut). Thousands of compounds are presently listed as HPPD inhibitors [15]; therefore, it is important to highlight that this differential toxicity depending on the mode of administration calls for a systematic comparison across a broad spectrum of inhibitors, as a way to reveal alternative compounds directed to the same enzyme.

The spreading of insecticide resistance among natural populations is a factor that has limited their efficiency in the control of vector-borne diseases and hence, fueled the search for alternative methods. Since mosquitoes can produce many generations per year, resistance can evolve very quickly. Besides, the appearance of cross-resistance among different neurotoxic compounds is a common finding [7,35,36]. A combination of different mechanisms, such as metabolic resistance, mutation of the target proteins and penetration factors (cuticular resistance) contribute to the resistance of insects to contact insecticides. Here we searched for cross-resistance using populations known to be resistant to organophosphates and pyrethroids, where both metabolic resistance (increased expression of detoxifying enzymes) and target-site mutations (*kdr*) are at play [24,25,27–30]. No evidence for cross-resistance to NTBC appeared using oral administration (Figure 4, Table 3), but when using the topical application assays a moderate (3.9 to 5.5 fold) but significant (p<0.001) increase in the LD50 was observed in Santarem, Nova Iguaçu and Oiapoque populations (Figure 5, Table 3). These results might be explained by a lower penetration of NTBC through the cuticle as a possible additional resistance mechanism present in these field populations, that complemented the role of metabolic resistance and target-site mutations. Several studies have reported increased expression of genes related to cuticle formation in resistant mosquito populations [37,38]. However, this does not fully explain the high levels of resistance to neurotoxic insecticides observed in these populations, pointing to a multifactorial nature of resistance.

Rock-kdr strain is derived from the backcrossing of a field-derived population (homozygous for *kdr* mutations) and the Rockefeller strain for eight generations, to reduce the contribution of detoxification enzymes (glutathione-S-transferase, esterases and multifunction oxidases) in pyrethroid resistance and to evaluate the effect of the kdr mutations alone [24]. As expected, since the molecular targets are different, when NTBC was orally or topically administered to the Rock-kdr strain, there was no significant variation in the LD50 with respect to the Rockefeller control strain. Similar results were observed when comparing the Red Eye strain and the Rockefeller strain (Figure 5, Table 3). Furthermore, NTBC was also lethal for *An. aquasalis* and *Cx. quinquefasciatus* with potency similar to that observed in *Ae. aegypti* (Red Eye and Rockefeller strains), reinforcing the hypothesis that it can be used to target several vector-borne diseases at the same time.

The effectiveness of topical application of NTBC suggests that it can be used in strategies such as indoor residual spraying (IRS) and Long-Lasting Insecticide-treated Nets (LLINs). However, HPPD inhibitors have a lethal effect only in blood-fed insects, a particular characteristic that makes them selective for hematophagous arthropods. This fact also creates some limitations for their use in topical-application strategies (such as LLIN and IRS), as illustrated by the results showing that the efficacy of NTBC in topical application trials strongly depended on the time interval between drug administration and blood meal intake. When mosquitoes were fed 24 hours after the topical application of an LD95 (the dose that killed 95% of mosquitoes when applied immediately after feeding), it only killed 50% of the insects, indicating that a significant proportion (around 50%) of the drug had already been inactivated or excreted by that time. NTBC LD95 only killed 25% of the mosquitoes when fed 72 hours after application, and it was not effective when mosquitoes were fed later on (Fig. 6). When NTBC was applied at different times after a blood meal (PBM), it was lethal when applied up to 48 hours PBM, indicating that most of the tyrosine had already been catabolized by that time (Fig. 7). Taken together, our data demonstrate that NTBC may be useful as a lead compound for developing compounds with a longer active life in mosquitoes. A wide range of other HPPD inhibitors have already been identified in the search for herbicides and should be investigated as possible tools for mosquito control. Additionally, the mortality determined by NTBC is not as fast as with the neurotoxic insecticides, and some fraction of a population of treated females may survive longer than the gonotrophic cycle, which takes around 2–4 days after the blood meal [39,40]. This fact may contribute to maintain susceptible alleles in the population, slowing the evolution of resistance. This would be an effect similar to that of the idealized late-acting insecticides [41].

The most used endectocide drug (it has activity against endo- and ectoparasites when applied to the host) in human and livestock is Ivermectin, capable of killing a wide variety of parasites and vectors [42,43]. It targets a broad range of parasites and invertebrates, including mosquitoes [44–47], and it has been proposed as an additional tool to control vector-borne diseases such as malaria [48–54]. However, ivermectin’s half-life in humans is short (around 18 h) and it would be necessary to administer multiple doses, a limitation in terms of logistics [52]. A recent study proposed the use of isoxazoline drugs (fluralaner and afoxolaner), currently used against fleas and ticks infesting animals, for the drug-based oral treatment of a proportion of human population for the control of *Ae. aegypti, Culex pipiens, Anopheles* and sand flies [55]. These compounds possess long *in-vivo* half-lives that provide weeks to months of protection after a single oral administration. However, they are not approved for use in humans and there is no information about their pharmacokinetic and/or possible side effects. In contrast, NTBC has been used in humans for treatment of HT-1 since 1994. It is remarkably safe drug for mammals (LD50>1000 mg/kg in rats) [56] presenting a half-life of 54 h in human plasma. Its concentration in blood following the ingestion of a therapeutic dose (1 mg/kg) is 8 μg/ml (24.3 µM) [57], much higher than the LD95 observed for all mosquito species during artificial feeding experiments. Because NTBC toxicity towards hematophagous arthropods is maximized when it is co-delivered along with the blood meal, our results raise the possibility that it could be used as a new endectocide drug (in humans and livestock) for control of mosquito-borne diseases, as part of an integrated vector management program.

## Acknowledgments

We wish to thank Dr. Martha Sorenson for a critical reading and for all suggestions in the writing of the manuscript. We thank all of the members of the Laboratório de Bioquímica de Artrópodes Hematófagos (UFRJ), especially JM Freire for breeding of *Ae. aegypti* (Red Eye strain) and J Marques, C Cosme and SR Cássia for technical assistance.

We also thank members of the Laboratório de Fisiologia e Controle de Artrópodes Vetores (FIOCRUZ), especially L dos Santos Dias, for suggestions for conducting the trials and for providing the resistant populations from Santarém and Nova Iguaçu, and L Carrara for providing Oiapoque population and Rock-kdr strain. We thank R Santos, P Serravale and Q Amorim for the breeding of *An. aquasalis* and *Cx. quinquefasciatus*.

